# nf-core/pairgenomealign: a scalable pipeline for pairwise whole-genome alignment

**DOI:** 10.64898/2026.11.18.767494

**Authors:** Charles Plessy, Mohammed Mahdi, Takahiro Fujita, Martin C. Frith, Nicholas M. Luscombe

## Abstract

Genome assemblies are accumulating rapidly across the Tree of Life, creating a growing need for reproducible and scalable methods for whole-genome comparison. We developed nf-core/pairgenomealign, a Nextflow workflow that automates pairwise genome alignment using the LAST software suite and produces standard alignment, visualisation, and summary outputs. The pipeline supports a wide range of evolutionary distances, runs reproducibly across local, cloud, and high-performance computing environments, and exports alignments in formats including MAF, BAM, BCF, and CRAM. We repurposed the compact format used for sequencing reads to combine the alignments of many genomes to the same target into a single file, which genome browsers display directly and from which multiple sequence alignments of any region can be extracted. Benchmarking against established genome aligners shows competitive performance with tunable trade-offs between speed and sensitivity, while applications to comparative genomics and assembly quality control illustrate its uses.

## Introduction

Recent advances in sequencing technologies, laboratory automation, and genome assembly algorithms have opened the prospect that chromosome-level genome assemblies may eventually become available for most known species. As genome assemblies become increasingly abundant, their comparison is emerging as a routine task for assembly quality control, variant discovery, evolutionary analysis, and large-scale comparative genomics. Although a rich ecosystem of specialised software exists for individual scientific questions, there remains a need for a generic workflow that aligns genome assemblies of diverse sizes, divergences and qualities, and delivers the alignments in standard formats for any downstream analysis, while prioritising ease of deployment, accessibility to non-specialists, reproducibility, robustness, and high-throughput execution. To address this need, we developed the nf-core/pairgenomealign pipeline using the Nextflow workflow system (Di Tommaso et al., 2017) and the community standards, reusable modules, and software-distribution ecosystem provided by nf-core (Ewels et al., 2020; Langer et al., 2025).

We used the LAST software suite (Frith et al., 2010) for pairwise genome alignment because it handles this diversity of lengths and divergences, as well as compositional biases, while implementing an efficient seed-and-extend strategy with successive gapless and gapped extension until similarity fades. The key advantages of LAST, which we leverage in this pipeline, are 1) its flex-ible seeding framework, which allows users to tune sensitivity and computational cost through the design of exact or inexact seeds, including spaced, transition-constrained, and sparsely sampled seeds (Noé and Kucherov, 2005; Frith et al., 2023), 2) its training procedure that automatically infers substitution scores and gap penalties from the data, enabling robust alignment across queries with heterogeneous evolutionary distances (Hamada et al., 2017), and 3) its last-split program, which selects an optimal set of non-overlapping alignments to produce a consistent one-to-one mapping between genomes, thereby resolving multi-mapping regions in a principled manner (Frith and Kawaguchi, 2015), which avoids the need for post hoc chaining and netting strategies (Kent et al., 2003).

Several alternative pairwise alignment tools were considered, including LASTZ (Harris, 2007), MUMmer4 (Marcais et al., 2018), Minimap2 (Li, 2018), and, more recently, FastGA (Myers et al., 2025). LASTZ and MUMmer4 are widely used for whole-genome comparison; LASTZ also offers configurable seeds and can infer scoring parameters from the data, but neither provides the probabilistic split-alignment framework of LAST. Minimap2 achieves excellent performance for read mapping, and its genome-alignment presets target closely related genomes. FastGA, which appeared after the core of this pipeline was designed, is much faster than the other aligners; we compare it with LAST in our benchmarks. We therefore selected LAST because it combined broad applicability across evolutionary distances with configurable sensitivity, data-driven parameter training, and principled resolution of ambiguous alignments.

Once computed, whole-genome alignments can be stored in formats that differ substantially in information content and storage efficiency. The Multiple Alignment Format (MAF), originally developed for the UCSC Genome Browser and related tools such as MULTIZ (Kent et al., 2002; Blanchette et al., 2004), is the primary output format of LAST. In addition to alignment coordinates, MAF stores the aligned sequences and, in LAST, per-base alignment confidence values. More compact formats trade information for storage efficiency. The tabular output of LAST and the PSL format used by BLAT (Kent, 2002) omit the alignment strings while retaining gap structure and summary statistics such as mismatch counts and gap openings. BLAST tabular output (Altschul et al., 1990) is more compact still, discarding the gap coordinates. At the end of the spectrum, alignments can be represented simply as paired genomic intervals in GFF or BED format with only an alignment score, providing near-maximal lossy compression in a text representation. Conversely, MAF alignments can be exported with minimal information loss to SAM/BAM (Li et al., 2009) and subsequently to CRAM format (Bonfield, 2022), benefiting from reference-based compression. The LAST suite can directly generate or export all of these formats.

The precursor of nf-core/pairgenomealign was developed for the genome alignment analyses of Plessy et al. (2024), including comparisons of extensively rearranged invertebrate genomes. We have since generalised, extended, and released this workflow as an nf-core pipeline. In this article, we describe its implementation, the improvements introduced since the original version, and its application to whole-genome comparisons across a broad range of evolutionary distances.

## Methods

Running the pipeline requires only a target genome, a list of query genomes and, optionally, the choice of a seed; all other settings have defaults suited to most comparisons. The rest of this section documents these defaults and the implementation, for users who need to adapt them.

### Software packaging, versions and documentation

The nf-core modules used in this pipeline rely on Free Software distributed through Bioconda (Gruening et al., 2018). Depending on the execution environment, these packages are installed automatically by Nextflow in isolated conda environments or provided through software containers. The default container sources are BioContainers (Leprevost et al., 2017; Bai et al., 2021), which maintain a one-to-one correspondence with Bioconda packages, and Wave containers (Seqera, 2026), which can combine packages from both Bioconda and conda-forge (community, 2015) within a single image. Containers are distributed in Docker-compatible formats (Di Tommaso et al., 2015) and can also be used through Singularity/Apptainer (Kurtzer et al., 2017), with BioContainers images converted and distributed by the Galaxy project (Afgan et al., 2016) and Wave images provided through Seqera. On shared computers, a common container cache lets first-time users run pipelines without waiting for software installation. Usage instructions (example command lines, sample-sheet format), the list of parameters with their defaults, and the description of the output files are maintained in the documentation of each version of the pipeline (for version 3.0.4: https://nf-co.re/pairgenomealign/3.0.4/docs/usage/, https://nf-co.re/pairgenomealign/3.0.4/parameters/ and https://nf-co.re/pairgenomealign/3.0.4/docs/output/), a copy of which is included in the source code archived on Zenodo (Plessy et al., 2026a).

The nf-core modules implementing the LAST alignment workflow use several minimal containers with partially overlapping software dependencies. To avoid repeated downloads of redundant images, the nf-core/pairgenomealign pipeline overrides the default module configuration and provides a single shared container containing LAST version 1652, samtools and bcftools (Danecek et al., 2021) version 1.24 with their library htslib (Bonfield et al., 2021), open-fonts 0.7.0 from conda-forge (required for rendering sequence names in dot plots), gzip 1.14, bzip2 1.0.8, and xz 5.8.3.

### Software configuration

Each module fills a command template: some tool arguments are hard-coded, and others come from generic variables (ext.args, ext.args2,…), through which users can add, change or remove arguments in a Nextflow configuration file. Which arguments are hard-coded can be seen in the source code of the module.

CPU cores, memory and run time are allocated to each process according to its nf-core resource label (from process_single to process_high), and can be changed in a Nextflow configuration file.

### Input

Input genomes are provided in FASTA format and may optionally be compressed with gzip. When downstream outputs are generated in BAM, BCF, or CRAM format, the pipeline ensures that the target genome is available in the block-gzip (BGZF) format by recompressing it when necessary (Li, 2011). Existing BGZF-compressed files are reused unchanged. The use of BGZF enables the random-access indexing required by these output formats. The target genome, which will be indexed, is specified with the pipeline parameter --target. We use *target* rather than *reference* because *reference genome* commonly refers to a species’ canonical genome assembly. Query genomes are specified through a sample sheet supplied via --input. As a convenience, a single query genome may alternatively be provided directly through the --query parameter.

### Alignment parameters

Prior to database construction, any existing lowercase soft masking in the target genome is discarded and replaced with tandem-repeat masking using the tantan implementation embedded in LAST (Frith, 2011). The maximal repeat unit length is hardcoded to 400 nt instead of lastdb’s default of 100, to accommodate primate centromeric regions. Mask replacement can be disabled with the pipeline parameter --softmask original, which preserves the masking present in the input assembly.

The target database is indexed on both strands using the lastdb option -S2. This increases memory usage during indexing and alignment but accelerates subsequent alignments. An important consequence is that some alignments may be reported between the reverse-complement strand of the target and the forward strand of the query, which software that assumes the target to be always on the forward strand will mishandle. Users may therefore revert to single-strand indexing with the pipeline parameter --strand forward. The default seed scheme is RY4 (sparse seeds that sample 1 in 4 positions; Frith et al., 2023); other seeds can be selected with the --seed parameter, such as the sparser RY128 or the more sensitive YASS (Noé and Kucherov, 2005) and MAM8 (Frith and Noé, 2014).

Alignment parameter training is performed with symmetric substitution scores and gap penalties by passing the --revsym, --gapsym, and --matsym arguments to last-train. These constraints are appropriate for most nuclear genomes but may be undesirable when evolutionary processes differ between strands, such as in animal mitochondrial genomes. In such cases, advanced users can override the training arguments by redefining ext.args for the ALIGNMENT_ TRAIN process in a Nextflow configuration file. Alternatively, a predefined LAST parameter file may be supplied through the pipeline parameter --lastal_params. The same parameter set is then applied to all query genomes. Such files may originate from a previous training run or correspond to a standard LAST scoring scheme.

By default, sequence alignments are computed with the lastal arguments -C2 -D1e9, configurable through the pipeline parameter --lastal_args. The -D1e9 setting specifies a significance threshold of one random alignment per 1 Gb of query sequence, whereas -C2 prunes nested alignments in order to reduce the computational burden associated with highly repetitive regions such as centromeres. During the reduction of many-to-many alignments to one-to-one relationships, alignments with a mismap probability (the probability of being aligned to the wrong locus) greater than 10*^−^*^5^ are discarded. This threshold is controlled by the pipeline parameter --last_split_mismap. We name these relationships by the multiplicity on the target, then on the query: in many-to-one (m2o) alignments, each base-pair of the query is aligned at most once, while base-pairs of the target can be aligned several times; one-to-many (o2m) alignments are the converse, and in one-to-one (o2o) alignments each base-pair of either genome is aligned at most once.

Multi-threaded LAST processes are allocated up to 12 CPU cores, subject to any lower limit imposed by user settings or institutional execution profiles. The number of threads does not affect the alignments, but with more than one thread, the order in which they are reported can vary between runs. The pipeline regression tests align SARS-CoV-2 sequences and compare the checksums of key output files with reference values.

### Export to SAM/BAM/BCF/CRAM formats

Alignments are converted from MAF to SAM with the maf-convert utility of LAST, and then to BAM, CRAM or BCF with samtools and bcftools, in an nf-core module that we extended for this pipeline. Although these formats were originally designed to store sequencing reads aligned to a reference genome, they can also represent whole-genome alignments. The pipeline exploits this capability by assigning a distinct read group to each query genome and encoding genome alignments as hard-clipped segments of the query sequence. In BAM and CRAM files, the length of each operation in the CIGAR string, which describes the alignment, is limited to 268,435,455 (2^28^ *−* 1) bases. When an alignment starts or ends further than this from an end of its query sequence, maf-convert writes the hard clip as several consecutive operations, so that alignments on query chromosomes and scaffolds longer than this limit are encoded without loss of information.

BCF files representing alignment mismatches as single-nucleotide polymorphisms are produced by passing -B to bcftools mpileup and --ploidy 1 -m to bcftools call. CRAM files are written in version 3.1 format by default. For compatibility with tools that do not read version 3.1, users may select a different CRAM version by modifying the ext.args2 setting of the ALIGNMENT_EXP process (one file per query) and the ext.args setting of the ALIGNMENT_MERGE process (combined file) in a Nextflow configuration file. Index files are generated automatically for all BAM, BCF, and CRAM outputs.

### Pairwise alignment plots

Dot plots of the alignments are drawn as bitmap images with the last-dotplot command. It defaults to a maximum of 100 sequences per genome, which guards against genome quality issues such as containing more than one million contigs. The target genome is plotted as a horizontal axis and the query as a vertical one, with the sequence names oriented so that they are readable in both cases. The target genome spans the width of the plot, set in pixels with the --dotplot_width option (default 1,200). The maximum height (--dotplot_height, default 10,000) is large enough that the width is effectively fixed, although the lengths of the query sequence names can cause small differences. Other options can be passed to last-dotplot with the --dotplot_options parameter. Finally, maf-linked can be used to denoise plots with the --dotplot_filter option. Plots of many-to-many alignments can be computationally expensive as well as uninformative when repeated regions are too frequent, and their generation can be turned off with the family of --skip_dotplot_… parameters. Runs of at least ten Ns in each assembly are drawn as pink lines, as they usually mark the boundaries between contigs in scaffolds.

### Statistics and metadata collection

Assembly statistics of the query genomes, including sequence length and base composition, are calculated with assembly-scan v1.0.0 (Petit III, 2023) unless disabled through the pipeline parameter --skip_assembly_qc. In addition, custom nf-core modules developed for this pipeline collect summary statistics from the alignment outputs and store them as text files within the results directory. These statistics include the nucleotide percent identity of the alignments computed with or without including gaps (May, 2004), gap frequencies, inferred insertion and extension penalties, and the nucleotide substitution matrix estimated from the whole-genome alignment. Coordinates of assembly gaps are identified with seqtk cutN v1.4-r122 (Li, 2025) using the arguments -n 10 -p 100000, to report runs of at least ten consecutive N bases. Optional thumbnail representations of one-to-one alignments can be generated with the pipeline parameter --multiqc_thumbs. Modules written for this pipeline then format the assembly statistics and thumbnails for a MultiQC report (Ewels et al., 2016), which consolidates them with the alignment statistics. The report of the full-size test of version 3.0.4 (human versus other primates) is included in the supplementary data (Plessy et al., 2026b) and displayed in the “AWS Results” tab of the pipeline’s website (nf-core, 2026).

### Hardware, datasets and benchmark procedures

All analyses were run with releases 3.0.2 to 3.0.4 of the pipeline, which run the same version of LAST (1652) with the same commands and therefore produce the same alignments. The pipeline was run with the Slurm scheduler (Jette and Wickberg, 2023) and Singularity on the high-performance computing (HPC) cluster of the Okinawa Institute of Science and Technology Graduate University (OIST), on nodes with Intel Xeon Gold 6230 CPUs (Figures 5 and 7) or AMD EPYC 7702 CPUs (comparison of CPU architectures), and by nf-core on Amazon Web Services nodes with Intel Xeon Platinum 8375C CPUs (full test). Myers et al. (2025) aligned the human genome CHM13 (GCF_009914755.1) to five mammalian genomes (human GRCh38 (GCF_ 000001405.40), chimpanzee (GCF_028858775.2), siamang (GCF_028878055.3), pig (Jinhua_ pig_T2T) and mouse (GCA_964188535.1)) on two Intel Xeon Gold 6226R CPUs, of the same generation (Cascade Lake) as the Xeon Gold 6230, with the same maximum clock frequency (3.9 GHz) and cache per core, and a higher base frequency (2.9 instead of 2.1 GHz). Nextflow reports the run time, memory use and CPU usage of each module run; to match the CPU time reported by Myers et al. (2025), we multiplied run time by CPU usage. For Figure 2, each of the 33 primate genomes was aligned to each of the others (1,056 ordered pairs) with the RY128 seed, to match the configuration of the pipeline’s full test suite, which runs on the Amazon Web Services cloud with a cap in computation resources that does not allow the use of the RY4 seed. The distance between two genomes was 100 minus the percent identity of their one-to-one alignment, counting gaps, averaged over the two alignment directions. Genomes were clustered with the average-linkage method (UPGMA).

**Figure 1.**
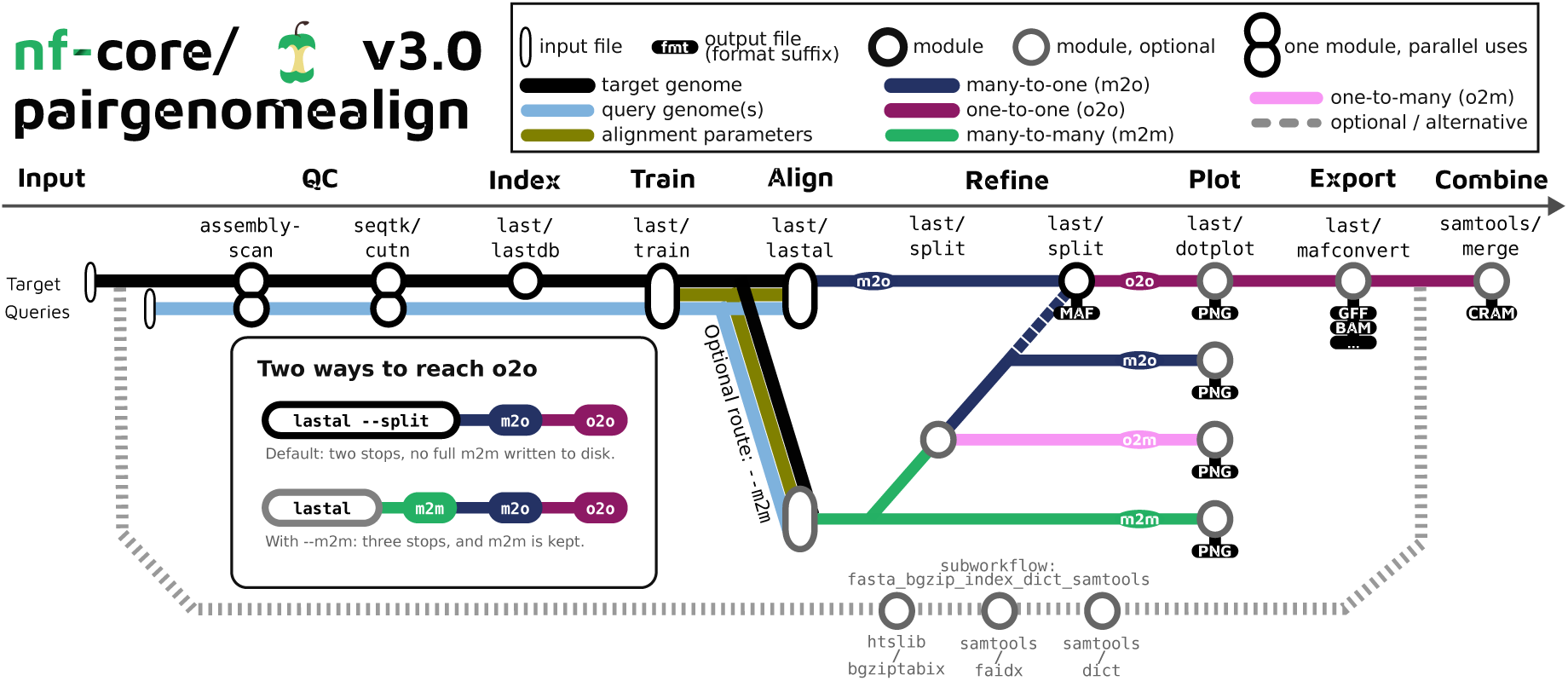
The nf-core/pairgenomealign pipeline aligns a list of query genomes to a target genome and produces alignments, plots, and metrics reports. The data flow is indicated by lines and the nf-core modules are indicated by circles. The main pipeline steps are indicated by words in bold. By default, the pipeline produces many-to-one alignments, where each region of the query genome is aligned at most once, and reduces them to one-to-one alignments, where each region of either genome is aligned at most once. The --m2m mode keeps the many-to-many alignments, where regions of both genomes can be aligned several times, and also produces one-to-many alignments, where each region of the target genome is aligned at most once.

**Figure 2.**
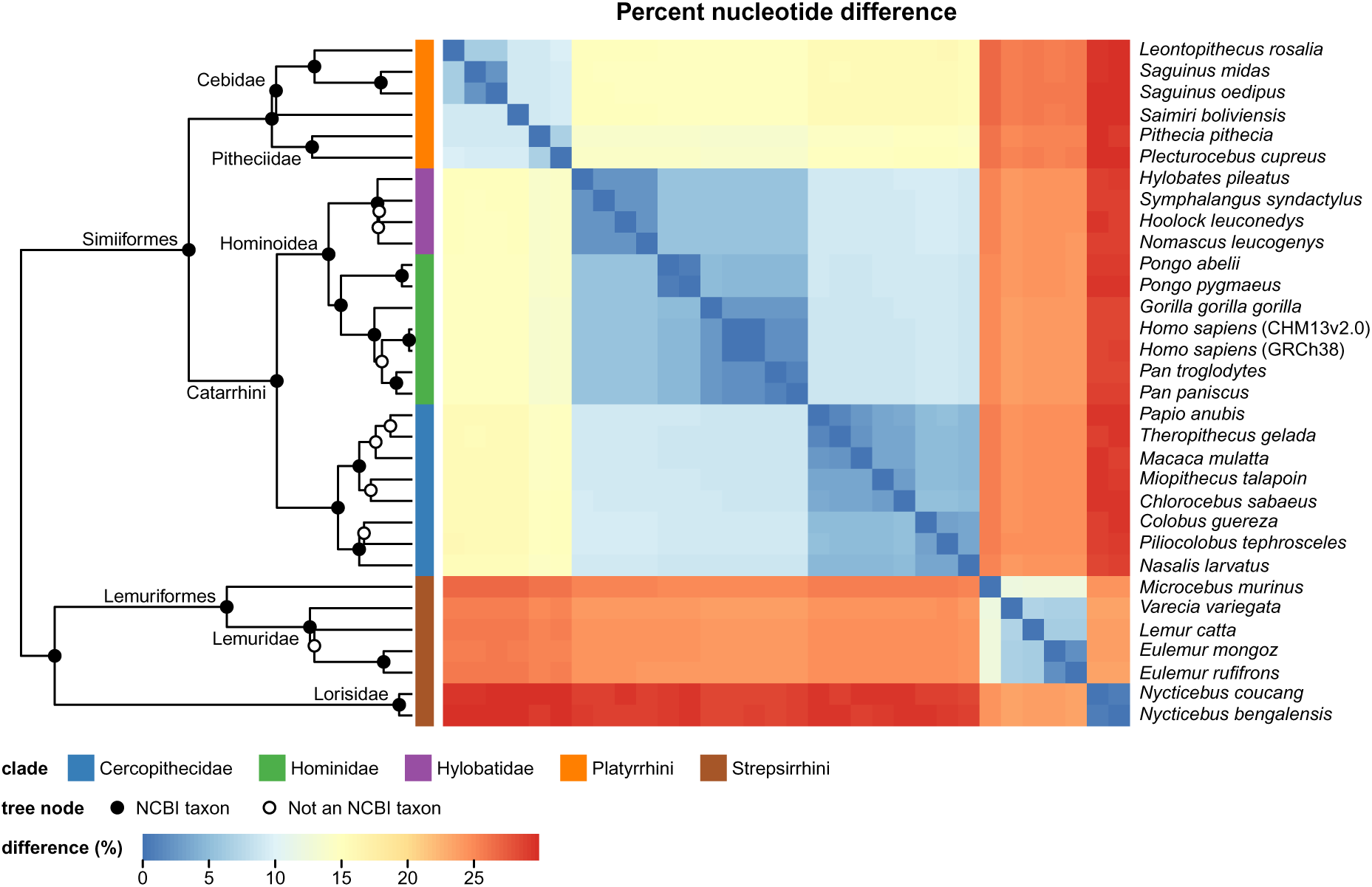
All-versus-all one-to-one alignment of primate genomes produces a matrix of percent nucleotide differences, plotted as a heatmap, which were clustered by average linkage; the tarsier is left out (see supplementary data). On the tree, filled dots mark the clades matching NCBI Taxonomy taxa (named when of family rank or above), and open circles the clades resolving a polytomy of the NCBI Taxonomy.

To repeat the coverage benchmark of Gulhan et al. (2025), we retrieved from the authors’ public Galaxy history (https://usegalaxy.org/u/cartman/h/fga-mm2-lz-last-comparison) 213,124 merged coding-exon intervals of the hg38 human genome, and the hg38 coordinates of their mm39 alignments made with KegAlign (a reimplementation of much of LASTZ accelerated on graphics processing units), LAST, FastGA and minimap2. We then ran nf-core/pairgenomealign 3.0.4 (LAST 1652) with default parameters in many-to-many alignment mode for each of the seeds RY128, RY4, YASS and MAM8, and counted the exons overlapped by an alignment and the coding bases covered on hg38. Running their LAST commands in the pipeline (LAST 1608; lastdb -uRY128; last-train --revsym -C2; lastal -C2; no last-split) overlapped the same exons as their LAST alignment. For the negative control, --reverse was passed to lastal with the --lastal_extr_args parameter, and the scores trained on the mouse genome with the --lastal_params parameter. The benchmark scripts are archived on Zenodo (Plessy et al., 2026b).

To repeat the simulation benchmark of Gulhan et al. (2025), we retrieved their 164 simulated sequence pairs and their alignments made with LASTZ v1.04.22 (default parameters), FastGA (commit 170a178) and minimap2 v2.28 (-x asm20), from their repository (https://github.com/nekrut/lz-fg-mm, commit ded50e5). We did not analyse the minimap2 alignments, because the published PAF files contain no base-level alignment: their coordinates are the ends of seed chains. Each pair consists of a random sequence of 1, 2, 5 or 10 kb (60% A+T) and a copy mutated with a substitution rate of 0 to 0.40, each flanked on both sides by 1 kb of independent random sequence. Although the repository describes indels at a rate of 1%, its simulation notebook inserts 1 to 5 bases in the copy at a rate of 0.005. We therefore computed the true identity of a pair as one minus the number of substitutions divided by the length of the homologous region. We aligned the pairs with the LAST commands and container image of nf-core/pairgenomealign 3.0.4 (LAST 1652), with the seeds RY128, RY4, YASS and MAM8. Trend lines are robust locally weighted regressions (Cleveland, 1979), computed with the R function loess (span 0.5, family = “symmetric”), and not drawn where fewer than 12 points lie within 3 percentage points of identity. Scripts are archived on Zenodo (Plessy et al., 2026b).

### Use of artificial intelligence

The pipeline was developed without AI agents that edit code or run commands; conversations with Microsoft Copilot and Seqera Co-Scientist were used for troubleshooting. For this article, the authors drafted the text as notes and paragraphs, which were edited with Microsoft Copilot. We then used Claude (Anthropic) as an agent with access to the data and the source code, to edit and draft parts of the text, to write the R and Python scripts that replicate the benchmarks, analyse their results and draw Figures 4 to 7, to rewrite the code that draws Figure 2, and to check statements of the manuscript against the source code of the pipeline. The authors checked these outputs and take responsibility for the content of the article. To check whether AI services propose a design similar to ours, we asked six of them in November 2026 to plan a pipeline that aligns query genomes to a target genome, with a short and a detailed prompt (prompts and answers are in the deposit). Two services proposed designs based on LAST, with alignment scores trained for each pair of genomes: Seqera AI, which named nf-core/pairgenomealign, and Claude, which did not name it but, in one answer, used its parameter and file names. The four other services proposed designs based on minimap2 or LASTZ, and none of them proposed to train the alignment scores for each pair of genomes.

## Results

### How it works

We developed the nf-core/pairgenomealign pipeline (Plessy et al., 2026a) to align in parallel a list of query genomes to a target genome. The main steps of the pipeline (Figure 1), described in detail in the methods section, are 1) index the target genome, 2) train the alignment parameters, 3) align each query to the target and 4) refine the alignments to one-to-one relationships where sequences of each genome are aligned at most once. The pipeline also collects assembly, alignment, and computational resource use statistics.

### Multiple genome alignment

From a list of genomes, all the alignment pairs can be produced by running the pipeline in a loop with each one playing the role of the target iteratively. We illustrate this by aligning 33 primate chromosome-level assemblies selected to maximally span the primate phylogenetic tree. The human versus other primates iteration corresponds to the *“full”* test suite of the pipeline, which took about 90 min to run on the Amazon computing cloud. In clades where the molecular clock is stable enough, hierarchical clustering of the percent nucleotide differences recovers trees consistent with the known taxonomy, as in the primates (Figure 2).

To combine the one-to-one alignments of all query genomes into a single indexed file, the pipeline repurposes the CRAM format, which was originally designed for read-to-genome representation but provides an efficient foundation for genome-to-reference representation too. In an all-versus-all alignment of the 33 primate genomes, the combined CRAM files were 5 to 9 times smaller than the compressed one-to-one MAF files of the same alignments. For any region, a multiple alignment projected on the target coordinates, leaving out insertions relative to the target, can be extracted efficiently, for instance with the pysam interface to HTSlib (Heger et al., 2026), and the aligned sequences, insertions included, can be passed to multiple sequence alignment software. Genome browsers display the file directly, which helps to inspect phylogenetic footprints. We illustrate it with an alignment of the human genome against the other 580 phase 1 assemblies of the Vertebrate Genomes Project (Formenti et al., 2026), including 13 invertebrate outgroups, which took 9 h on OIST’s HPC cluster and 490 CPU-hours. The resulting CRAM file takes 23 GiB of disk space, 4.5 times less than the compressed one-to-one MAF files of the same alignments (Figure 3).

**Figure 3.**
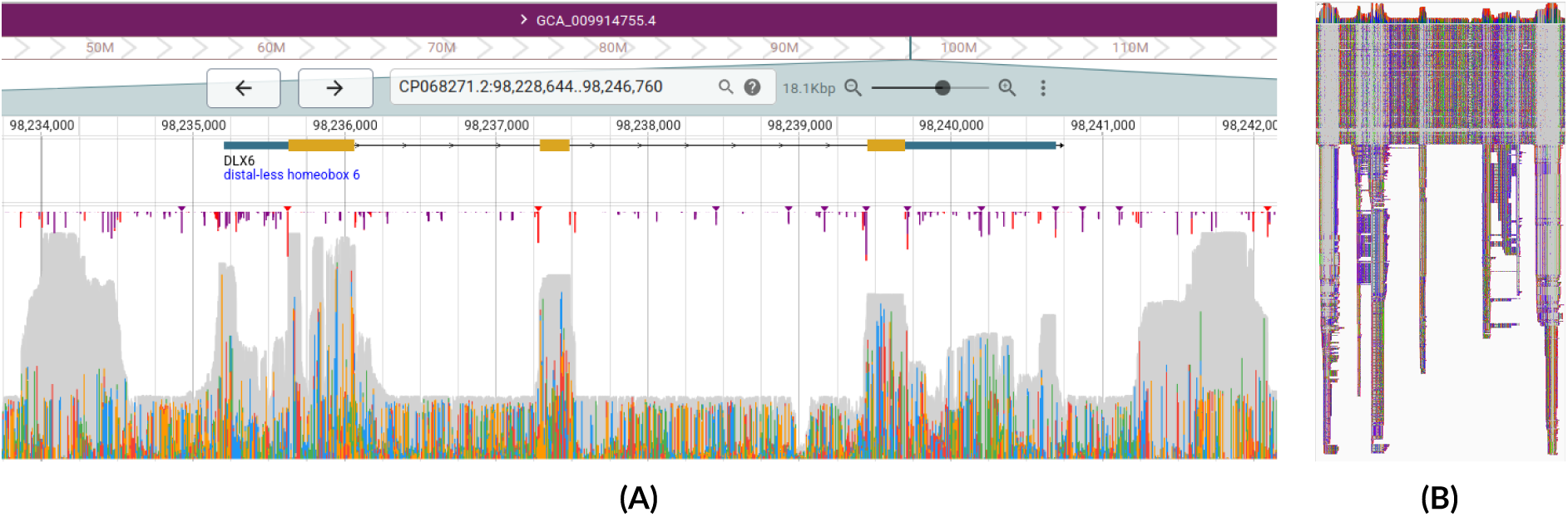
Human-versus-all one-to-one alignment of 580 genomes of the Vertebrate Genomes Project compressed in CRAM format displayed in genome browsers in the DLX6 locus, which is flanked by ultraconserved elements (UCEs, Bejerano et al., 2004). (A) Coverage track in the JBrowse genome browser (Diesh et al., 2023): its height is the number of genomes aligned at each position, in grey where their bases match the human genome and in colours where they differ; the grey peaks show where many genomes align without differences. (B) Alignment track of the same region in the Integrative Genomics Viewer (Robinson et al., 2011; Thorvaldsdottir et al., 2013), showing that all alignments can be displayed simultaneously.

### Performance and benchmark

To compare the sensitivity of the pipeline with that of other aligners where the true alignment is known, we used the 164 pairs of simulated sequences of Gulhan et al. (2025), in which homologous regions of 1 to 10 kb at 60 to 100% identity are flanked by random sequences (Figure 4A). We aligned them with the commands of the pipeline and compared the alignments with those published for the other tools. All the pairs were found by LASTZ and LAST with the YASS and MAM8 seeds; the default RY4 seed missed two pairs below 61% identity, the sparse RY128 seed found all pairs above 86% identity, and FastGA all pairs above 73% (Figure 4B). Training produced the same substitution and gap scores with RY4, YASS and MAM8, and where two of them found a sequence pair, the alignment ends were the same, except at 3 of the 324 ends. Seeds only determine where alignments start; from there, alignments are extended in both directions with the same scores. Where the alignment started from an RY4 seed stopped before the end of the homologous region, the denser YASS and MAM8 seeds started a second alignment beyond that point, which covered the rest. LASTZ alignments extended into the random flanks more often and further than LAST alignments (Figure 4B). Both aligners overestimated the identity of the most divergent pairs (Figure 4C). Identity is computed without gaps, so a gap that replaces mismatches by matches raises both the alignment score and the identity; LASTZ placed more gaps than the simulation had inserted. The precision of LAST’s alignment ends may come from its training of scoring parameters on the aligned sequences (Hamada et al., 2017): with parameters trained on the most divergent pairs only, its alignments of these pairs ended closer to the boundaries and overestimated identity less.

**Figure 4.**
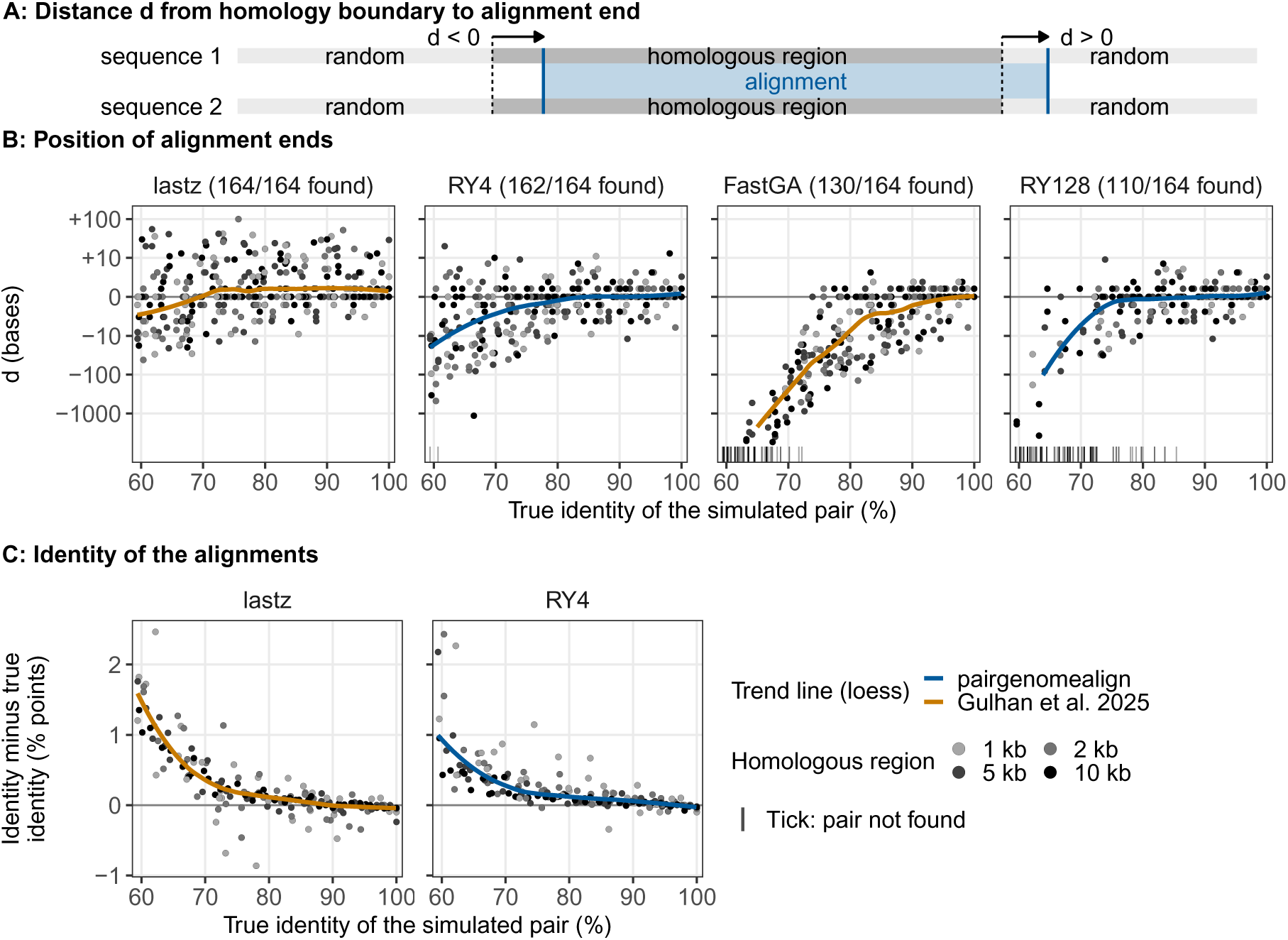
Alignment of simulated sequence pairs. (A) Each pair consists of a homologous region of 1 to 10 kb flanked by 1 kb of random sequence (Gulhan et al., 2025). *d* is the distance from the boundary of the homologous region to the end of the alignments, measured on the first sequence: negative when the alignments stop inside the homologous region, positive when they extend into random sequence. (B) *d* as a function of the true identity of the pair, for alignments published by Gulhan et al. (2025) (ochre) and for the many-to-many alignments of LAST run with the RY4 or RY128 seed (blue). Arc-sinh scale. Panel titles give the number of pairs found; ticks on the lower edge mark the pairs not found. YASS and MAM8 seeds (not shown) found all pairs, with nearly the same alignments as RY4. (C) Identity (without gaps) of the alignments minus true identity, in percentage points, for LASTZ and LAST (RY4). Grey levels: length of the homologous region. Lines: robust loess trends.

To compare computational cost and coverage on real genomes, we used the dataset of Myers et al. (2025), in which five mammalian genomes are aligned to a human genome. Using the same data, we ran our pipeline with three indexing seeds (RY128, RY4, and YASS) and measured (1) total CPU time, (2) peak memory usage, and (3) genomic coverage (Figure 5). Our CPUs were of the same generation as those of Myers et al. (2025) and at most as fast; on the logarithmic scale of Figure 5A, even a 2-fold difference in speed would change none of these comparisons except between nearly tied points. NUCmer and LASTZ completed the analysis within the memory available on most desktop computers, whereas FastGA required high-end consumer hardware. LAST, Minimap2, and Wfmash required workstation-, cloud-, or HPC-class resources. With a human genome as target, the pipeline needed up to about 40, 65 and 220 GiB of memory with the RY128, RY4 and YASS seeds, respectively (Figure 5A). LAST with the RY128 and RY4 seeds used less CPU time than all the other tools except FastGA (Figure 5A). Differences in coverage were largest for the most distant genome, mouse, for which coverage increased from FastGA to LAST with the RY128 and RY4 seeds, FastGA with gap filling, and LAST with the YASS seed (Figure 5B). Thus, gap filling restored the coverage of FastGA, but made it slower than LAST with the default RY4 seed, which covered nearly as much. On human coding sequences, the LASTZ (KegAlign) alignments of the mouse genome covered the most, followed closely by LAST with the MAM8 and YASS seeds (Figure 6). Coverage measures sensitivity only: alignments extending into non-homologous sequence also increase it, as LASTZ alignments did with the simulated pairs (Figure 4B). As a negative control, the mouse genome reversed but not complemented (Glidden-Handgis and Wheeler, 2024), which keeps its base composition and simple repeats but has no homology to the human genome, gave a single alignment of 178 bases with the default seed, and none in the one-to-one alignments. Overall, by changing a single parameter, the seed, the pipeline spans the range of speed and sensitivity covered here by several distinct tools.

**Figure 5.**
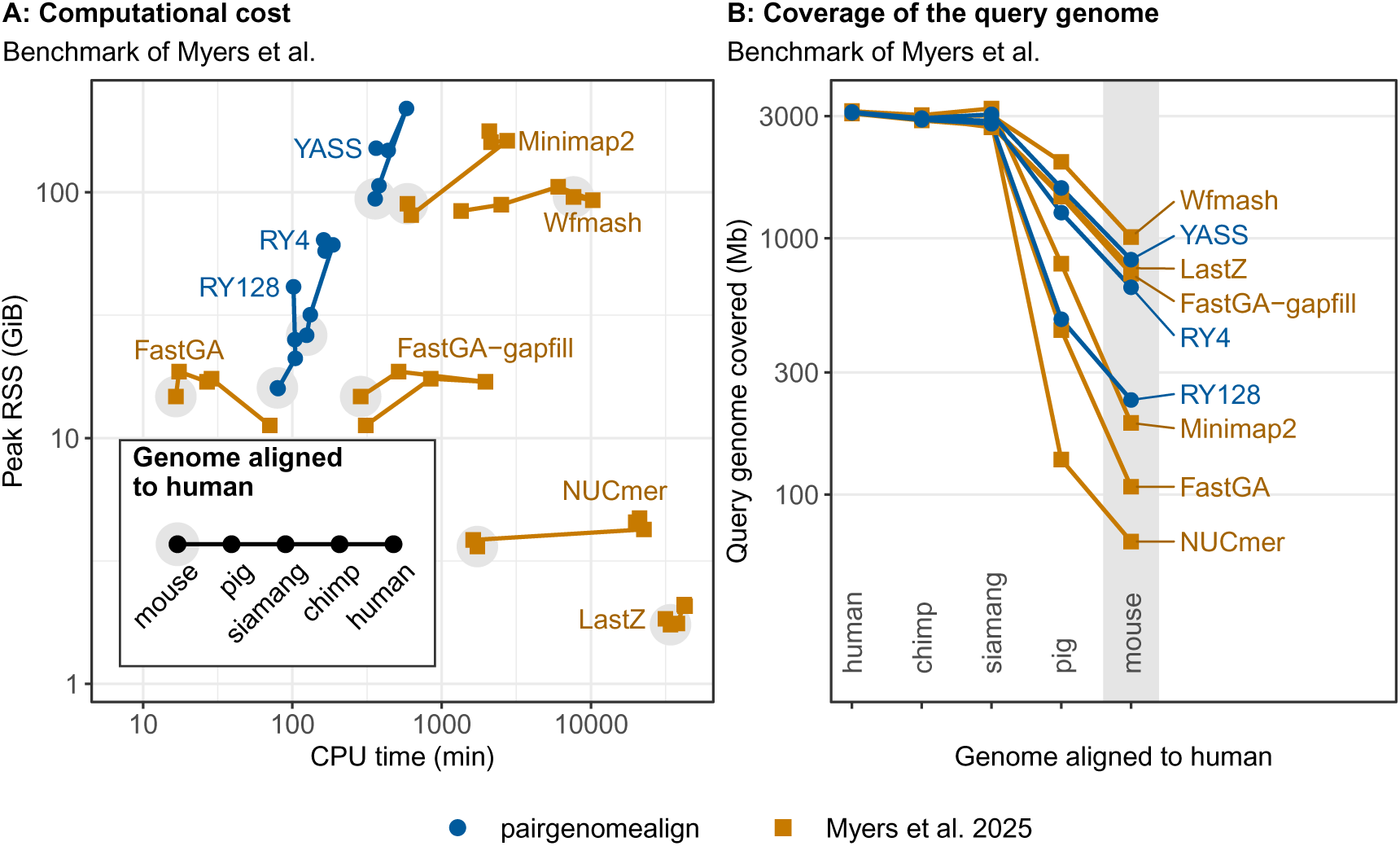
Pipeline performance with different seeds (blue) in the context of the benchmark of Myers et al. (2025) (ochre squares). Grey marks the mouse–human comparison in both panels. Human CHM13 genome aligned to five mammalian genomes including the human genome GRCh38. (A) Peak resident memory (RSS) versus total CPU time of indexing, training and alignment, on logarithmic scales. Comparisons made with the same software and parameters are connected in order of sequence divergence. (B) Length of the query genome covered by alignments (logarithmic scale); for pairgenomealign, many-to-one alignments.

**Figure 6.**
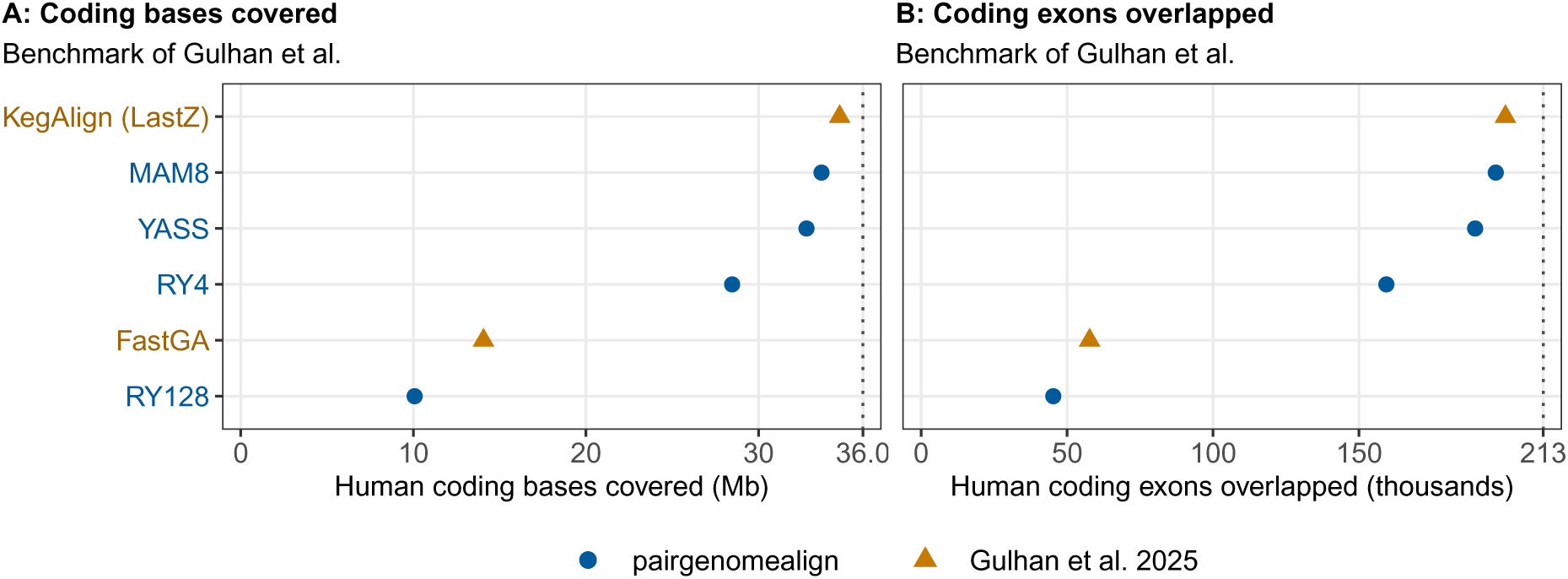
Pipeline performance with different seeds (blue) in the context of the benchmark of Gulhan et al. (2025) (ochre triangles). Mouse genome (mm39) aligned to the human genome (hg38). Human coding sequence covered by many-to-many alignments (--m2m mode), counted (A) in bases and (B) in exons overlapped by at least one aligned base. Dotted lines: complete coverage.

To detail how the pipeline uses computational resources, we ran each alignment three times on nodes with Intel Xeon Gold 6230 CPUs and three times on nodes with AMD EPYC 7702 CPUs, and quantified execution time and memory usage for indexing, training, alignment, and reduction to one-to-one relationships (“split”) (Figure 7). Alignment and splitting took most of the runtime, and alignment needed the most memory, regardless of seed choice. Indexing both strands of the human genome needed 10 to 30 GiB, depending on the seed. More distantly related genome pairs completed faster. Peak memory was the same on both CPU types, but on Intel nodes the wall-clock time was a median 19% shorter for alignment and 23% shorter for splitting.

**Figure 7.**
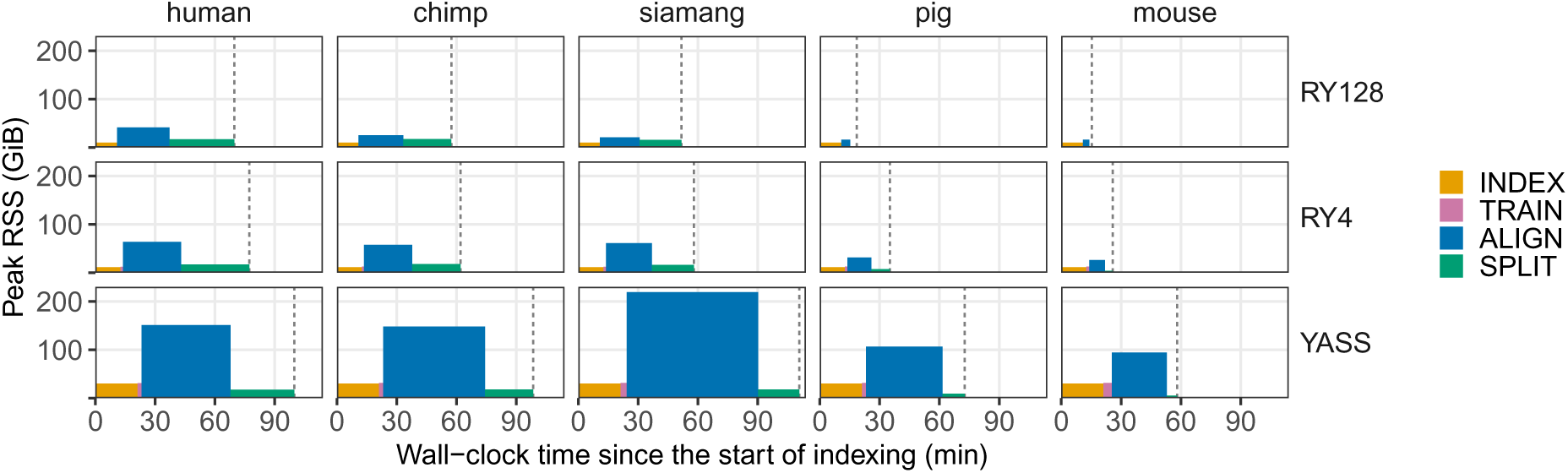
Wall-clock time and peak memory of each stage of the pipeline. Each panel shows the median for three replicate runs, aligning one of the five genomes of the benchmark of Myers et al. (2025) (Figure 5) to the human CHM13 genome with one seed on Intel Xeon nodes. Rows: seeds, in order of increasing sensitivity. Columns: aligned genomes, in order of increasing sequence divergence from human. Each stage is drawn as a block whose width is its wall-clock time and whose height is its peak resident memory (RSS). The dashed line marks the end of the run. In the default mode used here, ALIGN produces many-to-one alignments and SPLIT reduces them to one-to-one alignments.

These results support RY4 as the default seed. With simulated pairs, it missed only the most divergent ones and gave the same alignment ends as the denser YASS and MAM8 seeds; with the pig and mouse genomes, it covered about four fifths of the length covered by YASS, with about a third of its CPU time and memory.

### Pairwise plots for quality controls

Beyond evolutionary analyses, pairwise genome alignments are routinely used for assembly quality control. Draft assemblies containing unpurged or unresolved haplotypes often fail to exhibit a one-to-one correspondence with a reference genome. Similarly, diploid, polyploid, and whole-genome-duplicated genomes naturally produce non-unique alignments. The many-to-one or one-to-many dot plots generated by our pipeline, depending on which genome is the target, facilitate the detection of these features and provide an effective visual assessment of assembly quality. Homozygous regions appear as clear bands with no repeat-derived noise (Figure 8, where the target is a diploid assembly of *O. dioica* and the query one of its haplotypes).

**Figure 8.**
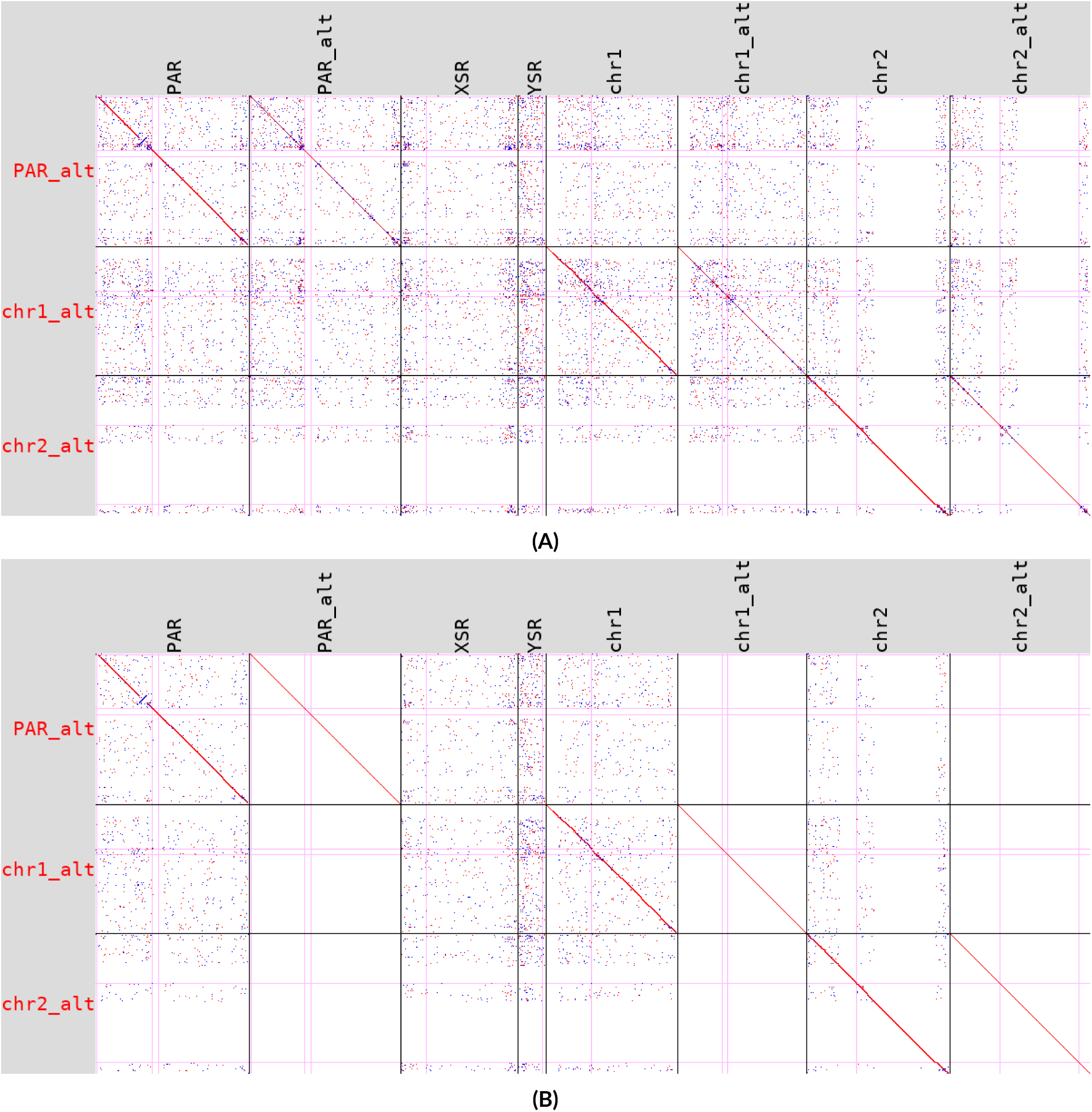
Alignment of one haplotype to the diploid assembly of a male *Oikopleura dioica* individual. Target (horizontal): diploid assembly, containing both haplotypes of each chromosome (suffix _alt for the alternative haplotype) and the X- and Y-specific regions (XSR, YSR) of the sex chromosome. Query (vertical): its alternative haplotype. Red and blue: alignments on the same and opposite strands; pink lines: contig boundaries within scaffolds. (A) Many-to-many alignment: repeats produce off-diagonal signal throughout. (B) One-to-many alignment, where each region of the target is aligned at most once: each query chromosome aligns to itself and to its homologue. In the columns of the alternative haplotype, the alignment of the query to itself covers every region, leaving no room for other alignments; off-diagonal signal remains only in target regions not covered by the homologous alignment, such as the Y-specific region, which has no counterpart in the query. Each pixel represents about 100 kb.

## Discussion

We developed a workflow that separates computationally expensive alignment generation from downstream interpretation, allowing users to revisit filtering, visualisation, and structural variation analyses without recomputing pairwise alignments. For instance, the one-to-one alignments can be loaded directly in the R package GenomicBreaks for synteny and breakpoint analyses, as in Plessy et al. (2024). By parallelising pairwise alignments, the pipeline scales naturally to all-versus-all genome comparisons and the construction of multiple sequence alignments. Whole-genome alignment workflows remain less common than alignment programs themselves. A widely adopted one is Cactus (Armstrong et al., 2020), which focuses on multiple-genome alignments, ancestral reconstruction, and pangenome representations, as does nf-core/pangenome (Heumos et al., 2024). Pairwise workflows include the chain-and-net pipelines of UCSC (Kent et al., 2003) and Ensembl Compara (Herrero et al., 2016) and make_lastz_chains (Kirilenko et al., 2023), all based on LASTZ, and nf-LO, which aligns genomes for lift-over with a choice of aligners (Talenti and Prendergast, 2021). nf-core/pairgenomealign targets reproducible pairwise genome comparisons using LAST, emphasising flexible sensitivity settings, standard genomics file formats, and straightforward deployment across local, cloud, and high-performance computing environments. By adopting the nf-core framework, the pipeline also benefits from community standards for documentation, testing, reproducibility, and longterm maintenance.

The main limitation of the pipeline is that it aligns genomes in pairs, to one target at a time: all-versus-all comparisons grow with the square of the number of genomes, and the combined CRAM file shows the other genomes only through the target. A single run, however, estimates the distances from the target to all queries, most accurately for the closest ones, so an iterative procedure could align only the pairs that an analysis needs. For phylogenetic footprinting, one run with a central target can suffice, such as the spotted gar for vertebrates (Braasch et al., 2016).

The workflow was designed to be customisable while retaining reproducibility. Users may replace the default container image, for example with institution-specific environments or other distributions of bioinformatics packages such as Debian Med (Moeller et al., 2010), pin a particular LAST version, or compile aligners with local hardware optimisations. Conversely, custom containers can preserve identical alignment software versions while benefiting from future developments of the workflow itself. The flexibility extends to execution strategies: while distributed execution through an HPC job scheduler is well suited to computationally intensive genome comparisons, analyses involving many small genome pairs may benefit from allocating a complete node and running the workflow locally within that allocation, thereby reducing scheduler overhead and improving resource sharing between tasks.

The pipeline also accepts sequencing reads in FASTQ format as queries. Parameter training then adapts the substitution and gap scores to the error profile of the sequencing technology, and the many-to-one alignments place each position of each read at most once on the target. While specialised read-mapping tools may be faster, using the same workflow for both genome and read analyses can provide practical advantages in large comparative projects.

The genomes aligned in this article span six orders of magnitude in size, from the 30-kb SARS-CoV-2 genomes of the regression tests to the 40-Gb genome of the West African lung-fish in the vertebrate comparison. We found the workflow to be tolerant of assembly and scaffolding quality. LAST can efficiently index and align highly fragmented genomes and unfiltered chromosome-level assemblies containing more than one million unplaced scaffolds (which unfortunately can be found in the databases of the International Nucleotide Sequence Database Collaboration (INSDC)). In practice, assemblies containing large numbers of unplaced scaffolds, unresolved haplotypes, or contaminant sequences can complicate interpretation and may expose limitations in downstream software, occasionally leading to excessive resource consumption or outright failures. While nf-core/pairgenomealign generally handles such datasets well, removing problematic sequences before analysis often improves interpretability and downstream robustness. This preprocessing step can be particularly beneficial when alignments are used for visualisation, synteny analysis, or structural variant discovery.

Several choices in nf-core/pairgenomealign depart from the most common practices and come from experience with comparative genomics. LAST provides the many-to-many, many-to-one, one-to-many and one-to-one alignments from a single tool, and its one-to-one alignments are a principled and efficient basis for comparing genomes at scale (Frith and Kawaguchi, 2015; Plessy et al., 2024). Storing genome alignments in CRAM, a format designed for sequencing reads aligned to chromosomes, allows the alignments of many genomes to the same target to be browsed together and sliced into multiple alignments; for this pipeline, the maf-convert utility of LAST was extended to encode alignments of the longest chromosomes. The masking of tandem repeats with units of up to 400 nt also reflects the use of the pipeline on large and repetitive genomes. Other features answer needs that a user asking only for alignments would not think of, but that make the results easier to interpret and check: the contig boundaries drawn on the dot plots, a MultiQC report summarising all the pairs, and the substitution matrix of each alignment. Finally, the pipeline is tested at each change of its code and on full-size genomes at each release, and, as noted above, handles low-quality assemblies, for instance containing millions of sequences.

### Conclusion

Our primary motivation in developing nf-core/pairgenomealign was to support genome-scale comparisons across the Tree of Life. In pursuit of this goal, we are applying the pipeline in ongoing comparative analyses, which will be reported separately. The workflow has been shaped by the practical demands of this undertaking, from handling imperfect public genome assemblies to efficiently exploiting modern computing environments. We anticipate that increasingly comprehensive catalogues of biodiversity will require reproducible and scalable approaches to pairwise genome comparison, and we hope that nf-core/pairgenomealign will provide a useful foundation for these future efforts.

## Acknowledgements

We thank the DNA Sequencing Section and the Scientific Computing and Data Analysis Section of the Research Support Division at OIST for their support; Priscila Biller, Simon Hellemans, Roger Huerlimann and Nicholas Treen for critical comments on the manuscript; Christopher Cheshire and the Luscombe laboratory for the Nextflow hackathons; Jiashun Miao and Huyen Pham for testing the pipeline on vertebrate genomes; and the nf-core community for reviewing the releases of the pipeline and running its full-size tests on Amazon Web Services.

## Data, script, code, and supplementary information availability

The nf-core/pairgenomealign pipeline is available at https://nf-co.re/pairgenomealign, and version 3.0.4, used in this manuscript, is archived on Zenodo (https://doi.org/10.5281/zenodo.22119145; Plessy et al., 2026a).

Scripts and data supporting this article are archived on Zenodo (https://doi.org/10.5281/zenodo.24327550; Plessy et al., 2026b): the scripts that run the benchmarks and draw the figures, with a README; the tables derived from the pipeline outputs and execution traces behind Figures 2 and 4 to 7; the MultiQC report of the full-size test of version 3.0.4; the prompts given to AI services and their answers; and the accession lists of the 33 primate assemblies and of the 581 phase 1 assemblies of the Vertebrate Genomes Project.

